# Dissolved oxygen minimally affects magnetic susceptibility in biologically relevant conditions

**DOI:** 10.1101/2021.03.13.434266

**Authors:** Véronique Fortier, Ives R. Levesque

## Abstract

**Purpose:** To investigate the potential of quantitative susceptibility mapping (QSM) with MRI as a biomarker for tissue oxygenation in fat-water mixture. Oxygen molecules (O_2_) are paramagnetic. This suggests that dissolved O_2_ in tissue should affect the measured magnetic susceptibility. However, direct measurements of dissolved O_2_ in tissues is challenging with QSM as the induced change in susceptibility is below the sensitivity of existing algorithms. QSM in regions that contain fat could be sensitive enough to be used as a marker of tissue oxygenation as oxygen has a larger solubility in fat than in water.

**Methods:** The relationship between dissolved O_2_ concentration and magnetic susceptibility was investigated based on MRI measurements using phantoms made of fat-water emulsions. Dairy cream was used to approximate fat-containing biological tissues. Phantoms based on dairy cream with 35 % fat were designed with controlled concentrations of dissolved O_2_. O_2_ was bubbled into the dairy cream to reach O_2_ concentrations above the concentration at atmospheric pressure, while nitrogen was bubbled in cream to obtain O_2_ concentrations below atmospheric pressure. Magnetic susceptibility was expected to increase, becoming more paramagnetic, as O_2_ concentration was increased.

**Results:** Magnetic susceptibility from MRI-based QSM measurements did not reveal a dependence on O_2_ concentration in fat-water mixture phantoms. The relationship between susceptibility and O_2_ was weak and inconsistent among the various phantom experiments.

**Conclusion:** QSM in fat-water mixture appears to be minimally sensitive to dissolved O_2_ based on phantom experiments. This suggests that QSM is not likely to be sensitive enough to be proposed as a marker for tissue oxygenation, as the change in magnetic susceptibility induced by the change in dissolved O_2_ concentration is below the current detection limit, even in the presence of fat.

## Introduction

Tissue oxygenation (pO_2_) mapping with magnetic resonance imaging (MRI), also referred to as MR-oximetry, is of high interest in the context of oncology, where the reduced presence of oxygen (O_2_) in tumors is known to be linked to resistance to cancer treatments [1], [2]. MR-oximetry is also of interest in the context of metabolic diseases, such as diabetes, as the lack of O_2_ in adipose tissue has been shown to lead to adipose tissue inflammation, which can result in insulin resistance [3].

Quantitative susceptibility mapping (QSM) can be used to measure O_2_ saturation in blood, thanks to the oxygenation-state-dependent magnetism of the hemoglobin molecule [4]. Oxyhemoglobin, the oxygenated form of hemoglobin, is diamagnetic (Δχ_oxy_ = −0.11 ppm relative to water at 22 °C [5], [6]), very similar to human soft tissue [7]. On the other hand, deoxyhemoglobin is paramagnetic (Δχ_deoxy_ = 3.46 ppm relative to water at 22 °C [5], [6]), and therefore induces positive contrast in QSM [7]. In healthy individuals, arterial blood is close to saturated and nearly all hemoglobin molecules carry O_2_ in arterial blood (normal value = 96 to 98% [8]). The proportion of oxyhemoglobin is reduced in venous blood (normal value = 70 to 80% [9]), due to the consumption of O_2_ in tissues. The relationship between the magnetic susceptibility and the deoxyhemoglobin concentration is linear [10], [11]. The slope of the blood magnetic susceptibility normalized to the hematocrit as a function of the deoxyhemoglobin concentration was shown to be equal to the difference in magnetic susceptibility between fully oxygenated and deoxygenated red blood cells, 3.57 ppm [5], [6], [12]. One can therefore derive blood O_2_ saturation directly from a QSM measurement, as previously proposed notably for brain [13] and heart [14].

Tumor oxygenation has been previously studied using QSM. Those studies were performed using QSM in combination with a hyperoxic gas challenge, where the participant breathes a gas mixture with a large concentration of O_2_ (more than 95 % O_2_) for a specific period of time. This was achieved in patients with brain tumors [15], and in a mouse model of colorectal liver metastases [16]. Blood oxygenation measured by QSM was used in both studies as a potential biomarker of tumor oxygenation. In both studies, variations of the mean magnetic susceptibility in a tumor region-of-interest (ROI) were observed while breathing hyperoxic gas, compared to breathing normal medical air. These observations of baseline magnetic susceptibility differences between healthy tissues and tumors, and of susceptibility variations in response to a hyperoxic gas challenge, were encouraging for the use of QSM as a biomarker in the context of MR-oximetry, keeping in mind that blood oxygenation is indirectly related to tissue oxygenation. Further investigation is needed to assess the potential of QSM for direct characterization of tissue O_2_ concentration.

O_2_ molecules are paramagnetic and can be dissolved in tissue interstitial fluid. This suggests that QSM might also be directly sensitive to tissue oxygenation. A change in the concentration of dissolved O_2_ molecules in tissue interstitial fluid could potentially be measured directly with QSM.

The sensitivity of QSM to tissue oxygenation can be investigated with a two-compartment model. This model could be used to describe the global magnetic susceptibility in a volume of tissue, combining information from blood and tissue interstitial fluid, as previously proposed [12], [17]. This model can be hypothesized to enable the simultaneous characterization of blood and tissue oxygenation with QSM. The model assumes that the bulk magnetic susceptibility of tissue is the sum of contributions from tissue and blood. The model is described by *Eq. 1*, where Δχ is the global magnetic susceptibility measured with QSM, V_Blood_ is the relative blood volume, Δχ_Blood_ is the magnetic susceptibility of blood, and Δχ_Tissue_ is the bulk magnetic susceptibility of the tissue, which is the parameter of interest for mapping of tissue oxygenation.

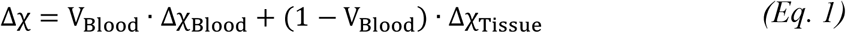

It is also assumed that the two contributions to the magnetic susceptibility are sensitive to the presence of O_2_: in blood via the oxygenation state of hemoglobin and the presence of dissolved O_2_ in the blood plasma, and in tissue via O_2_ dissolved in tissues interstitial fluid. Δχ_Blood_ can be modeled with *Eq. 2*, where Hct is the blood hematocrit, OEF is the oxygen extraction fraction, which corresponds to the relative portion of O_2_ removed by the tissue from the blood as it flows through the capillary bed [18], Δχ_oxy_ is the magnetic susceptibility of oxyhemoglobin, Δχ_deoxy_ is the magnetic susceptibility of deoxyhemoglobin, and Δχ_plasma_ is the magnetic susceptibility of the blood plasma. The values of Δχ_oxy_, Δχ_deoxy_ and Δχ_plasma_ are reported in the literature [5], [6]. Hct can be measured using a blood sample. OEF and V_Blood_ can be both measured with quantitative MRI techniques such as the quantitative blood oxygen level-dependent method (QBOLD) [19].

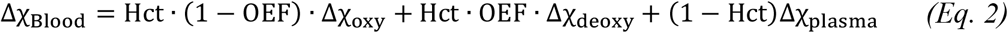

The change in magnetic susceptibility (Δχ) of a fluid induced by a change in the concentration of dissolved O_2_ molecules is small. A recent study demonstrated that the Δχ induced by a variation of oxygenation from normal pO_2_ (∼ 150 mm Hg) to near complete O_2_ saturation, (pO_2_ ∼ 650 mm Hg) is on the order of 30 ppb (0.03 ppm) [5]. These measurements were performed with a technique called MR susceptometry, which is a simplified but powerful approach to estimate the magnetic susceptibility that is valid when the geometry of interest can be approximated by an infinite cylinder [4]. That study demonstrated that a large change in O_2_ concentration in water induced only a very small, but significant, variation in magnetic susceptibility, with a measured slope of 0.062 ± 0.002 ppb/mmHg O_2_. A Δχ change of 30 ppb is at or below the measurement sensitivity of current in vivo QSM techniques. This suggests that QSM-based magnetic susceptibility measurements may not be sensitive enough to detect changes of dissolved O_2_ in tissue.

Despite its limited sensitivity to dissolved O_2_ in water, QSM could be a marker of dissolved O_2_ in fat-water mixtures, based on the fact that O_2_ is six times more soluble in fat than in water [20]. Prior work did not study the potential role of a fat component [5]. In this work, it was hypothesized that QSM may be sensitive to dissolved molecular O_2_ in fat-water mixtures [15]. This hypothesis was investigated using QSM measurements performed in a variety of fat-water phantoms made of dairy products.

## Methods

### Identification of the optimal dairy product for phantom design

Initial experiments were performed to identify the most appropriate dairy solution to use in this work to mimic in vivo fat-water mixtures at different O_2_ concentrations. Commercially available dairy products with different fat content (milk with 2% fat, and cream with 10, 22.5 and 35 %) were investigated. The homogeneity of the fat-water emulsion at room temperature was evaluated, as well as the stability of the dissolved O_2_ content over time. The ideal solution should feature high homogeneity of the fat and water phases and a good capacity to retain dissolved O_2_ concentration (i.e. a stable concentration of dissolved O_2_ over time).

The various dairy products were placed in closed 50 ml centrifuge and first exposed to room temperature for four hours. The tubes were filled with 42.5 ml of product, leaving an air space, and sealed with special screw caps with a self-sealing cap membrane (SeptaSecure Uncut Cap, Syringa Lab Supplies Inc.). Thread seal tape was used to improve the seal of the screw cap. After four hours of storage at room temperature, O_2_ was added to saturate the solutions. This was done by bubbling the solution with O_2_ for 10 minutes using a needle inserted through the self-sealing membrane, connected to a controlled O_2_ supply via plastic tubing. A second needle was also inserted for air to flow out of the tube during bubbling. Six vials of each solution with different fat content were prepared, for a total of 24 vials. An O_2_ flow rate of approximately 1 L/min was used.

The O_2_ concentration, the temperature, and the atmospheric pressure were measured in each vial using a dissolved O_2_ meter with an optical pO_2_ probe (Orion Star A323, Thermo Fisher Scientific). The probe was calibrated in air according to the manufacturer’s instructions. The absolute uncertainty of the probe was calculated based on the set of measurements described in Supporting information (*Supporting Table S1*). The O_2_ measurements were performed once in each vial, after six different waiting times (0, 15, 30, 60, 120, and 240 minutes). The characterization of O_2_ concentration measurements with the probe in dairy cream compared to distilled water is presented in *Supporting Figure S1*.

### Magnetic susceptibility phantoms

QSM experiments were performed in various fat-water phantoms to evaluate the sensitivity of magnetic susceptibility to O_2_ variations in the presence of fat. All phantoms included vials filled with dairy cream with 35 % fat with a range of O_2_ concentrations. O_2_ concentrations were obtained by bubbling O_2_ or nitrogen (N_2_) for a specific period of time, using the bubbling technique described in the previous subsection. A gas pressure of 51.7 kPa was used for N_2_ gas bubbling. All vials were placed in a larger container filled with distilled water to enable the measurements of non-local phase variations around the vials introduced by the differences in magnetic susceptibility [21], [22]. Gadolinium-based (Gd^3+^) contrast agent was added to the water compartment to reduce the T_1_ of water. The design of phantom #1 consisted of seven 50 ml vials filled with 42.5 ml of dairy cream at different concentrations of O_2_, and is shown in *Figure 1*. The gas bubbling times used for this phantom are presented in *Table 1*.

**Figure 1:**
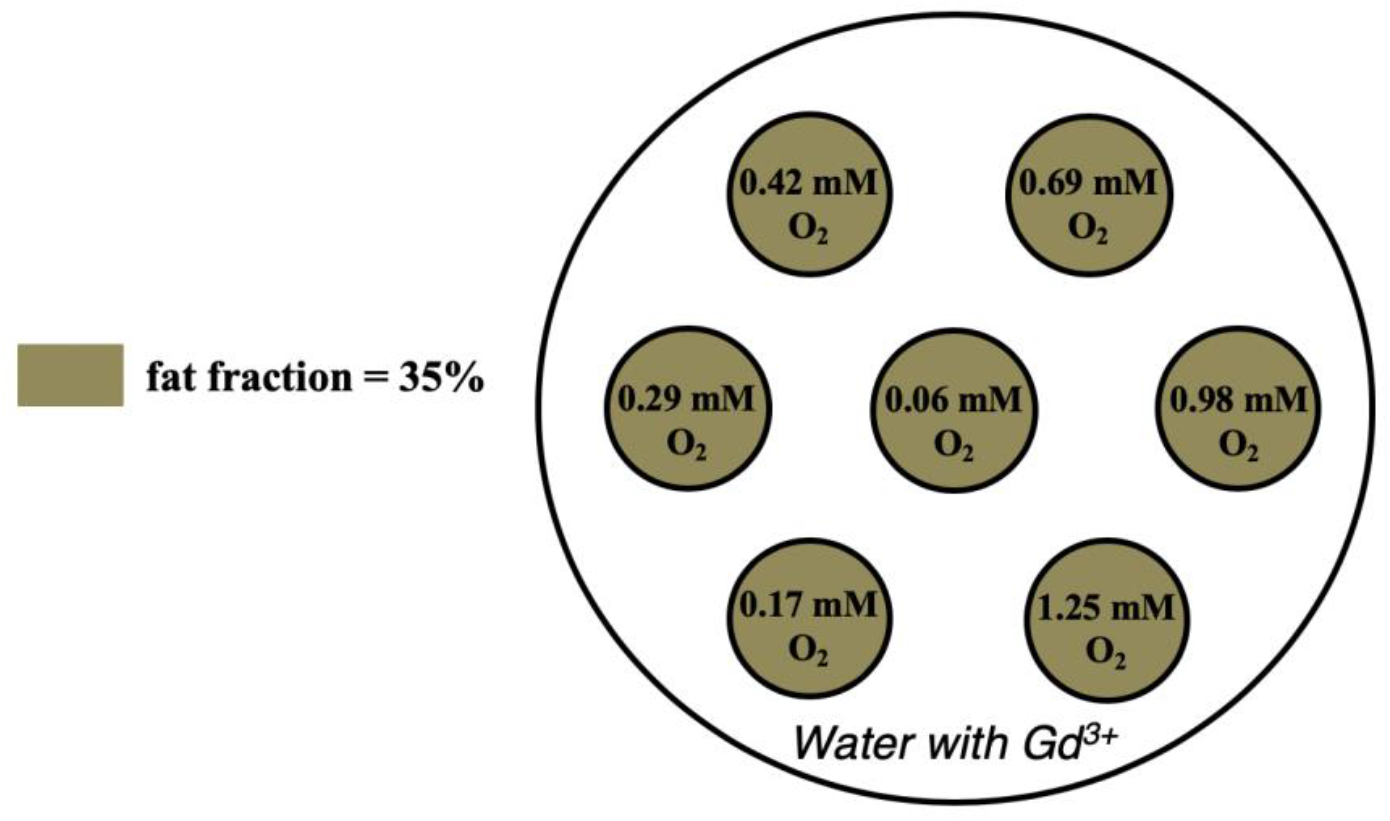
Phantom #1 based on dairy cream with 35 % fat. The *O*_2_ concentration was controlled in each vial. The *O*_2_ concentrations in the figure were the targeted values. The exact *O*_2_ concentration was measured in each vial with a probe and are reported in the Results section.

**Table 1:**
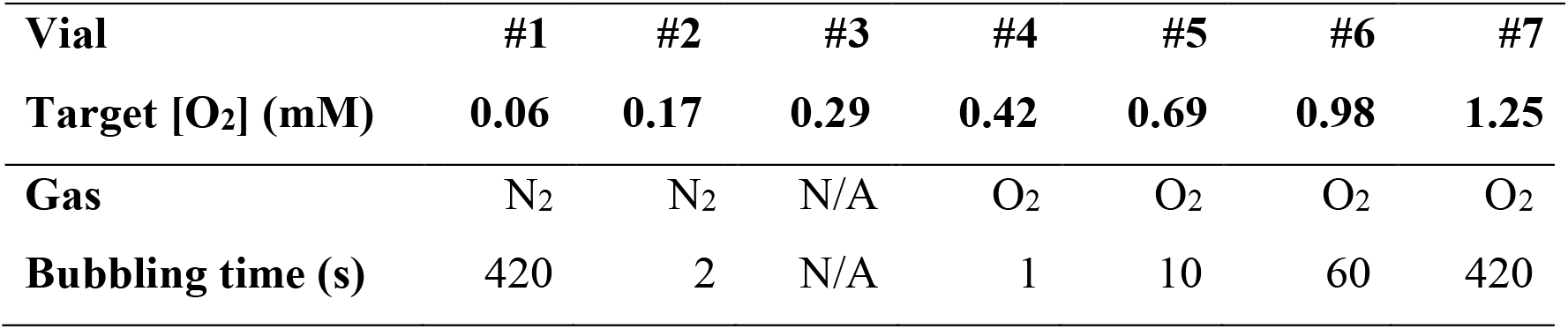
Gas and bubbling times used in phantom #1

| **Vial** | **#1** | **#2** | **#3** | **#4** | **#5** | **#6** | **#7** |
| --- | --- | --- | --- | --- | --- | --- | --- |
| **Target [O<sub>2</sub>] (mM)** | **0.06** | **0.17** | **0.29** | **0.42** | **0.69** | **0.98** | **1.25** |
| **Gas** | N<sub>2</sub> | N<sub>2</sub> | N/A | O<sub>2</sub> | O<sub>2</sub> | O<sub>2</sub> | O<sub>2</sub> |
| **Bubbling time (s)** | 420 | 2 | N/A | 1 | 10 | 60 | 420 |

Phantom #2 was designed based on phantom #1, but without N_2_ bubbling. This was done to remove the potential confounding effect introduced by the difference in magnetic susceptibility between N_2_ molecules, which are diamagnetic [23], and O_2_ molecules, which are paramagnetic [24]. In our observation, it is possible that the change in magnetic susceptibility observed in phantom #1 was due to a variation in the concentration of N_2_, instead of O_2_, or to a combination of both. Removing N_2_ bubbling avoids this potential confounding factor. Phantom #2 is shown in *Figure 2*. The gas bubbling times are presented in *Table 2*.

**Figure 2:**
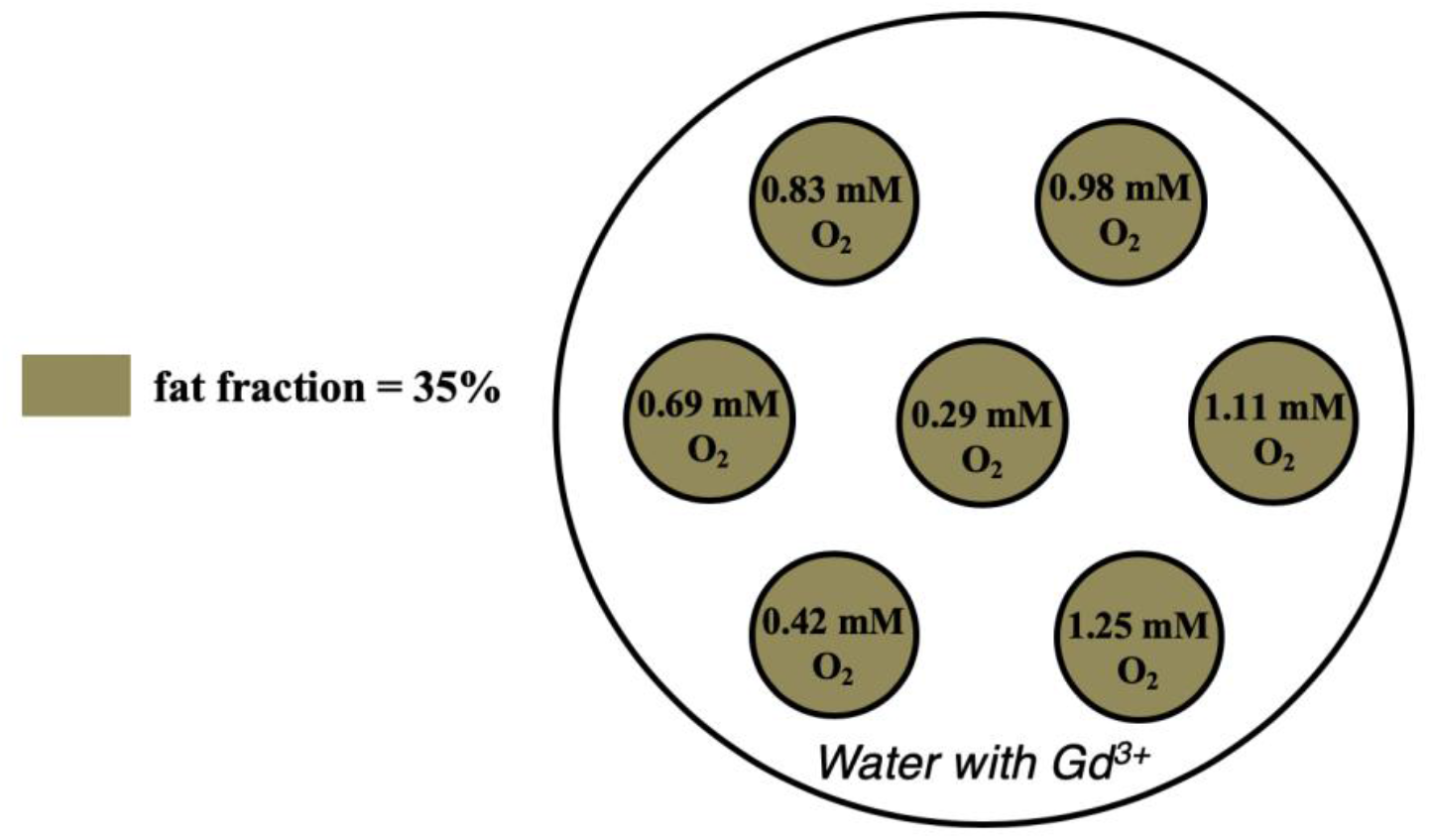
Phantom #2 based on dairy cream with 35% fat. The *O*_2_ concentration was controlled in each vial, and the figure shows the targeted values. The exact *O*_2_ concentration was measured in each vial with a probe and reported in the Results.

**Table 2:** Gas and bubbling times used in phantoms #2 and #3

| **Vial** | **#1** | **#2** | **#3** | **#4** | **#5** | **#6** | **#7** |
| --- | --- | --- | --- | --- | --- | --- | --- |
| **Target [O<sub>2</sub>] (mM)** | **0.29** | **0.42** | **0.69** | **0.83** | **0.98** | **1.11** | **1.25** |
| **Gas** | N/A | O<sub>2</sub> | O<sub>2</sub> | O<sub>2</sub> | O<sub>2</sub> | O<sub>2</sub> | O<sub>2</sub> |
| **Bubbling time (s)** | N/A | 1 | 2 | 5 | 10 | 30 | 420 |

Phantom #3 was used to replicate a prior phantom design used for susceptometry measurements [5]. Phantom #3 contained a single 15 ml vial filled with dairy cream in which the O_2_ concentration was set for each measurement, as illustrated in *Figure 3*. In this case, the phantom was imaged multiple times, changing the inner vial each time to cover all O_2_ concentrations. The targeted O_2_ concentrations were the same as in phantom #2, with only O_2_ bubbling and no N_2_ bubbling. The gas bubbling times are presented in *Table 2*. O_2_ bubbling was performed in a 50 ml vial filled with 42.5 ml of dairy cream, as described earlier, and 15 ml of the dairy cream was then transferred in the 15 ml vial using a syringe, through a needle inserted in the self-sealing membrane of the cap. A second needle was also inserted in the membrane to allow gas to flow out of the vial while transferring the dairy cream. Phantom #3 was prepared twice, once to perform QSM and MR susceptometry measurements with short TR and TEs, and once to perform MR susceptometry with long TEs.

**Figure 3:**
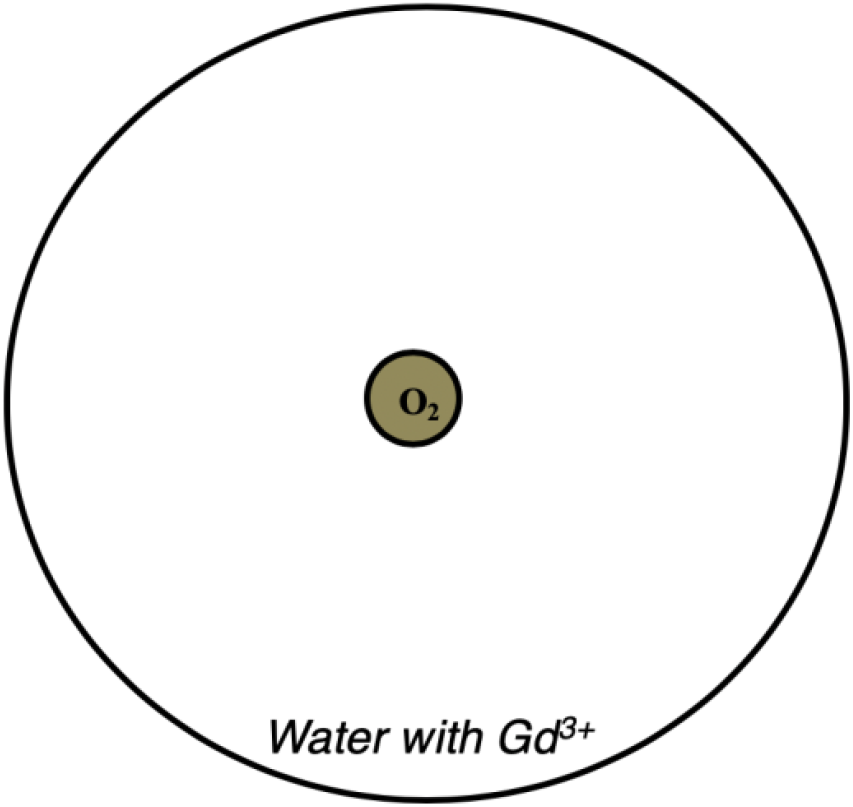
Phantom #3 based on dairy cream with 35% fat. The dairy cream vial had a specific concentration of *O*_2_, which was controlled by *O*_2_ bubbling.

Phantom #4, very similar to phantom #1 but with six 50 ml vials instead of seven, was designed to investigate the impact of the use of longer TEs on the relationship between the QSM-based magnetic susceptibility measurements and the O_2_ concentration. Longer TEs, up to 69.8 ms, have been proposed in the literature in the context of QSM-based MR oximetry measurements [15]. Such long TEs are expected to enable more phase contrast accumulation over time, which could be used to capture the expected relationship between QSM and O_2_ concentration. Gd^3+^-based contrast agent was added to the dairy cream in this phantom at a concentration of 0.10 mM to reduce the large T_1_ difference between the fat and water components to a value closer to what is observed in vivo. The design of phantom #4 is shown in *Figure 4* and the gas bubbling times used for this phantom are presented in *Table 3*.

**Figure 4:**
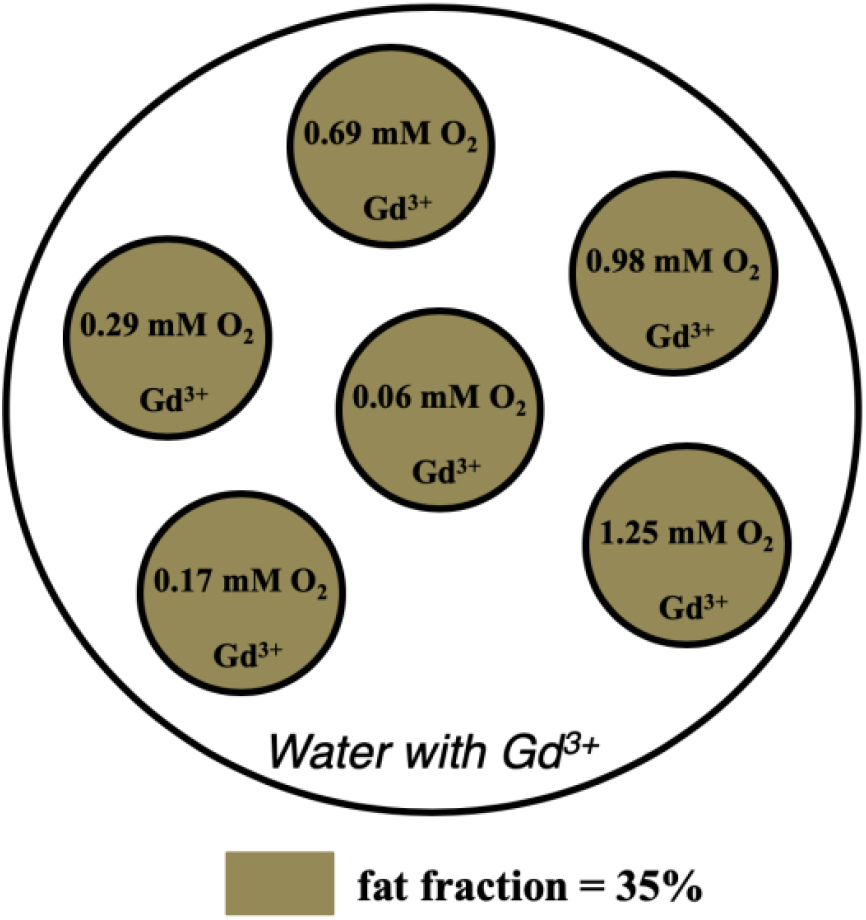
Phantom #4 based on dairy cream with 35 % fat. The *O*_2_ concentration was controlled in each vial. The *O*_2_ concentrations in the figure were the targeted values. The exact *O*_2_ concentration was measured in each vial with a probe and is reported in the Results.

**Table 3:** Gas and bubbling times used in phantom #4

| **Vial** | **#1** | **#2** | **#3** | **#4** | **#5** | **#6** |
| --- | --- | --- | --- | --- | --- | --- |
| **Target [O<sub>2</sub>] (mM)** | **0.06** | **0.17** | **0.29** | **0.69** | **0.98** | **1.25** |
| **Gas** | N<sub>2</sub> | N<sub>2</sub> | N/A | O<sub>2</sub> | O<sub>2</sub> | O<sub>2</sub> |
| **Bubbling time (s)** | 420 | 2 | N/A | 1 | 20 | 420 |

### Data acquisition

The data acquisition was performed with a 3 T MRI system (Ingenia, Philips Healthcare). A head coil was used for imaging of all phantoms. For phantoms #1, #2, and #4, the dairy cream vials were positioned perpendicular to the main field axis, while they were positioned parallel to the main field axis for phantom #3. A 3D spoiled multi-echo gradient-echo (MGRE) sequence was used with a monopolar readout gradient in all phantoms for QSM measurements. Two MGRE measurements were collected in each phantom with this pulse sequence, each with a different first TE. The measurements were then combined, alternating TEs from each MGRE measurement. A short first TE and TE spacing (ΔTE) were selected to enable accurate fat-water separation. Isotropic voxels with a large coverage in the slice orientation were used to maximize QSM accuracy, based on [25]. The sequence parameters are listed for all experiments in *Table 4*. 2D MGRE MR susceptometry data were also acquired in phantom # 3 based on the technique used in prior work [5].

**Table 4:** Imaging parameters for the multi-echo gradient echo sequences

|  | **Phantom #1** | **Phantom #2** | **Phantom #3** |  |  | **Phantom #4** |
| --- | --- | --- | --- | --- | --- | --- |
|  | **QSM** | **QSM** | **QSM** | **Susceptometry** |  | **QSM** |
|  |  |  | **Exp. #1** | **Exp. #1** | **Exp. #2** |  |
| Repetition time (ms) | 18 | 18 | 18 | 18 | 176 | 176 |
| Slice orientation | Coronal | Coronal | Axial | Axial | Axial | Coronal |
| Number of averages | 4 | 4 | 4 | 8 | 4 | 1 |
| In plane voxel size (mm<sup>2</sup>) | 1.15×1.15 | 1.15×1.15 | 1.3×1.3 | 1.15×1.15 | 1.15×1.15 | 1.15×1.15 |
| Slice thickness (mm) | 1.15 | 1.15 | 1.3 | 5 | 5 | 1.15 |
| Number of slices | 50 | 50 | 20 | 1 | 1 | 25 |
| TE<sub>1</sub> (ms) | I: 1.5 | I: 1.5 | I: 1.5 | I: 1.5 | I: 1.5 | I: 1.5 |
|  | II: 2.7 | II: 2.7 | II: 2.7 | II: 2.7 | II: 2.7 | II: 2.7 |
| ΔTE (ms) | 2.4 | 2.4 | 2.4 | 2.4 | 2.4 | 2.4 |
| Number of TEs | 3 | 3 | 3 | 3 | 28 | 28 |
| Flip angle (°) | 15 | 15 | 15 | 15 | 15 | 15 |
| Bandwidth (Hz/pixel) | 1041 | 1041 | 1181 | 1033 | 1029 | 1033 |

The QSM data processing in the presence of fat requires fat-water separation, which in turn requires a fat spectrum. Rather than using a generic fat spectrum from the literature, a measured spectrum of dairy cream was obtained using MR spectroscopy (MRS). A single voxel stimulated echo acquisition mode (STEAM) MRS acquisition was used, with a voxel size of 12×12×12 mm^3^, the minimum mixing time (TM = 16 ms), the minimum TE (10 ms), a repetition time of 6 s, and 16 signal averages. STEAM MRS measurements were performed once for each phantom, in the vial at room O_2_ concentration.

The O_2_ concentration, the temperature, and the atmospheric pressure were measured in each vial after MRI data acquisition using a dissolved O_2_ meter with an optical pO_2_ probe (Orion Star A323, Thermo Fisher Scientific), following calibration in air performed according to the manufacturer’s instructions.

### o_2_ measurements

Oxygenation measurements in g/L ([O_2_]_meas_) obtained with the probe were converted to mM units using *Eq. 3*, where 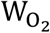 is molar mass of O_2_ (= 31.999 g/mol).

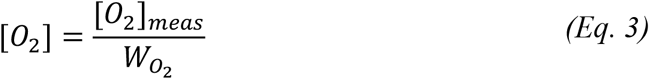

### Data processing

All data analysis and calculations were performed using software implemented in MATLAB (R2016b, The MathWorks, Natick, MA).

The raw MRS data were processed using custom software. Processing steps included a zeroth-order phase correction, followed by a Gaussian filter apodization in the time domain, and a Fourier transform. The resulting spectra were fitted to a nine-resonance spectral model (eight fat resonances and the water resonances) using least-squares fitting. The relative abundance of each fat resonance was calculated as the ratio of the area of each fat resonance relative to the total area of all fat resonances.

For QSM in all phantoms, fat-water separation was first performed. This was done using a two-step combination of the 3-Point Dixon algorithm [26] and the iterative decomposition of water and fat with echo asymmetry and least squares estimation (IDEAL) [27], where the output of 3-Point Dixon was used as the input to IDEAL, and the output of IDEAL was taken as the final result. First, the MGRE datasets (ΔTE = 2.4 ms) were combined by alternating the echoes, resulting in a dataset with a ΔTE of 1.2 ms. 3-Point Dixon was then performed on the complex data from the shortest TEs (1.5, 2.7, and 3.9 ms) of the combined dataset. The multi-resonance fat spectrum with eight fat resonances obtained from the single voxel STEAM MRS measurement was used. The 3-Point Dixon algorithm yields a B_0_ variation field map that ideally is free of chemical shift effects. This B_0_ map was then used as an initial guess for the IDEAL algorithm. The complex data from all six echoes combined were used for IDEAL to improve the accuracy of the fat-water separation and of the final B_0_ field map free of chemical shift effects.

The B_0_ field variation map obtained from fat-water separation was nominally free of chemical shift effects and was used as the starting point for QSM data processing. Unwrapping was performed on the field map using a quality-guided (QG) region-growing approach to remove any remaining spatial phase wraps in the 3D volume [28], after first scaling the data by the echo spacing to obtain a phase difference image. The background field effects were removed using the Laplacian boundary value (LBV) algorithm [29]. This step nominally removes all field effects from spatial variations in the static field of the MRI system and ideally leaves only the field variations due to the phantom composition. Using this processed field map, the magnetic susceptibility maps were reconstructed using the morphology enabled dipole inversion (MEDI) algorithm, a regularized least-squares approach to inverting the dipole field equation [30]. Finally, all magnetic susceptibility maps were referenced to the magnetic susceptibility measured in the dairy cream vial at O_2_ concentration under atmospheric pressure (nominal O_2_ concentration of 0.29 mM).

For MR-susceptometry data processing, the B_0_ field variation map free of chemical shift effects obtained from fat-water separation was also used as the starting point. No spatial unwrapping was required in this case due to the absence of spatial phase wraps in the 2D slice. The background field effects were removed using the regularized-enabled sophisticated harmonic artifact reduction for phase data (RESHARP) [31], as this is appropriate for 2D imaging data. The magnetic susceptibility was then calculated using *Eq. 4*, where f_0_ is the central frequency and Δf_0_ is the mean frequency shift measured in a manually drawn ROI within the cylindrical vial [5]. The uncertainty of the magnetic susceptibility obtained with *Eq. 4* was calculated based on error propagation, using the standard deviation of the frequency shift within the ROI as the uncertainty on Δf_0_. Magnetic susceptibility measurements were referenced to the magnetic susceptibility measured in the dairy cream vial at O_2_ concentration under atmospheric pressure (nominal O_2_ concentration of 0.29 mM).

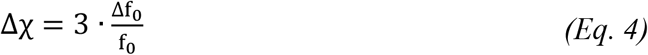

The correlation between magnetic susceptibility and O_2_ concentration was investigated in all datasets. The average and standard deviation of the magnetic susceptibility measured with QSM were calculated in 2D circular ROIs manually drawn in each vial on the middle slice of each dataset. Linear fits were performed between the mean magnetic susceptibility values respectively measured in all vials with QSM and MR susceptometry, and the measured O_2_ concentration.

## Results

### Identification of the optimal dairy product for phantom design

Dairy cream with 35 % fat saturated with O_2_ was the only dairy product that showed a stable O_2_ concentration after four hours when stored in a sealed container at room temperature, starting from an initial fully saturated state. This is illustrated in *Figure 5*. The O_2_ concentration in milk with 2 % fat decreased quickly as a function of the waiting time. This decrease in O_2_ concentration was slower in dairy products with increasing fat content. Dairy cream with 35 % fat was therefore used for all subsequent experiments.

**Figure 5:**
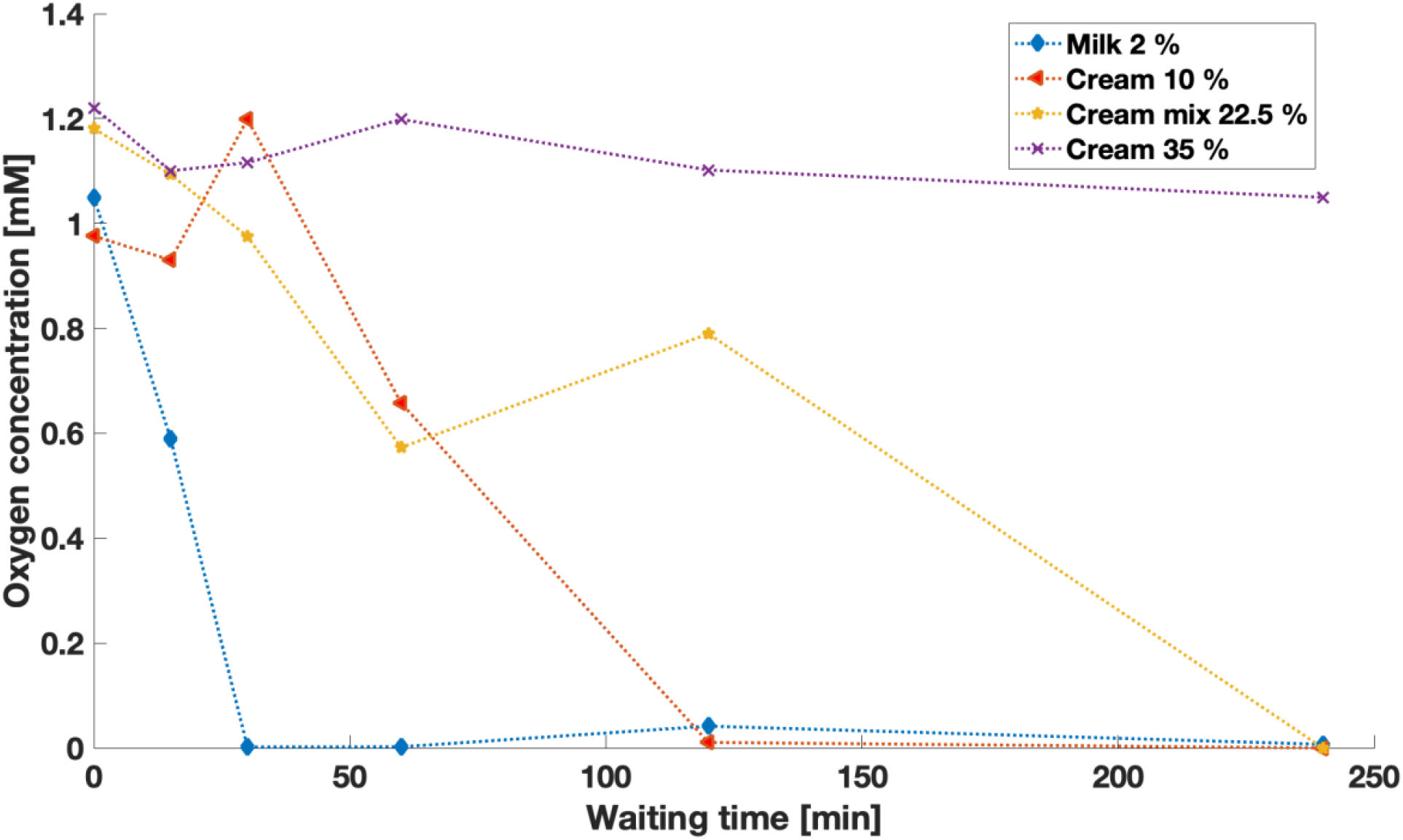
Oxygen concentration measured in various dairy products as a function of the waiting time after 10 minutes of bubbling with *O*_2_.

### O_2_ measurements

The O_2_ concentration in all vials of the four oxygenation test phantoms measured with the dissolved O_2_ probe are presented in *Table 5*.

**Table 5:** *O*_2_ concentration measured in all test phantom vials, in mM. The O _2_ probe has an absolute uncertainty of 0.01 mM.

|  | **Vial:** | **#1** | **#2** | **#3** | **#4** | **#5** | **#6** | **#7** |
| --- | --- | --- | --- | --- | --- | --- | --- | --- |
| **[O<sub>2</sub>]**<br>**(mM)** | Phantom #1 | 0.03 | 0.09 | 0.28 | 0.96 | 1.13 | 1.07 | 1.10 |
|  | Phantom #2 | 0.27 | 0.51 | 0.70 | 0.82 | 0.88 | 0.88 | 0.55 |
|  | Phantom #3, Exp. 1 | 0.26 | 0.31 | 0.59 | 0.83 | 0.75 | 0.48 | 1.01 |
|  | Phantom #3, Exp. 2 | 0.29 | 0.34 | 0.38 | 0.51 | 0.52 | 0.66 | 0.57 |
|  | Phantom #4 | 0.09 | 0.21 | 0.27 | 0.35 | 0.96 | 1.05 | - |

### Magnetic susceptibility phantoms

The relationship between magnetic susceptibility and O_2_ concentration observed in phantom #1 was moderate. A weak coefficient of determination (R^2^) was obtained for the linear fit (R^2^=0.42), with a positive slope. The results are shown in *Figure 6*.

**Figure 6:**
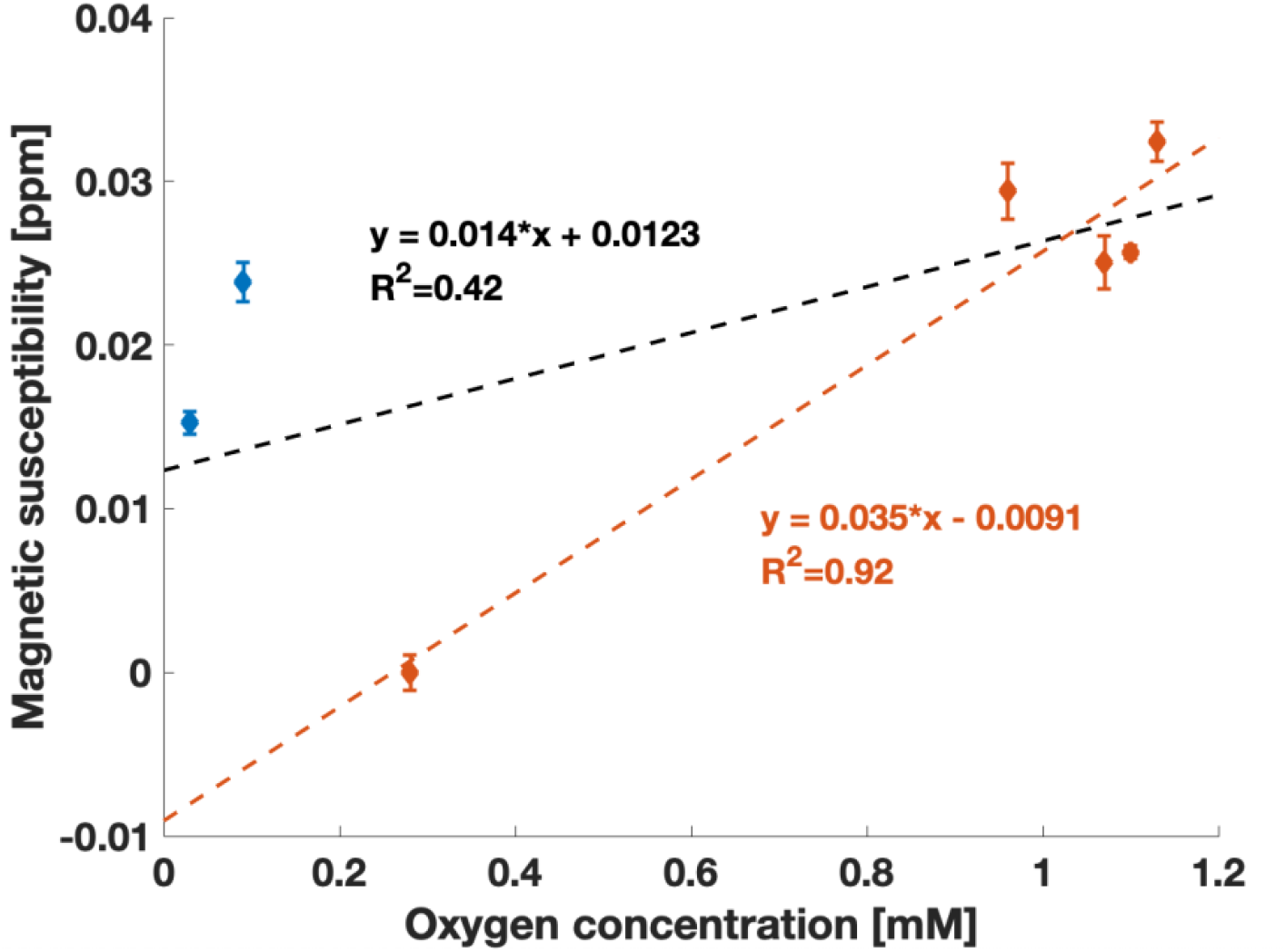
Variations of the magnetic susceptibility as a function of the *O*_2_ concentration measured in phantom #1. The mean (± standard deviation) magnetic susceptibility measured in a 2D ROI in each vial as a function of the measured *O*_2_ concentration is plotted. The orange points correspond to vials bubbled with *O*_2_ and the blue points to the vials bubbled with *N*_2_. The black dashed line corresponds to the linear fit over all points, while the orange dashed line corresponds to the linear fit over the orange points only. The vial at room *O*_2_ concentration (nominal *O*_2_ concentration of 0.29 mM) was used as the QSM reference (to 0 ppm).

N_2_ was used to replace O_2_ in two vials of phantom #1 to reduce the O_2_ concentration below atmospheric pressure, but this seemed to affect the magnetic susceptibility measurements in a way that does not fit with the expected variation as a function of O_2_. In this phantom, the two points corresponding to N_2_ bubbling did not follow the same trend as the points corresponding to O_2_ bubbling. A significant positive correlation with a R^2^ of 0.95 was obtained when the datapoints bubbled with N_2_ were ignored. However, this trend was mostly due to the widespread difference between the non-bubbled vial and the vials bubbled with O_2_. In addition, in this phantom, the two points corresponding to N_2_ bubbling showed a more paramagnetic susceptibility compared to the point at O_2_ concentration under atmospheric pressure, while these would be expected to be more diamagnetic. Based on these observations, N_2_ was removed from the design of phantoms #2 and #3 and the focus was shifted to O_2_ concentrations above atmospheric pressure (> 0.29 mM).

In phantoms #2 and #3, the magnetic susceptibility measured with QSM did not show any trends with O_2_ concentration. The results obtained with phantom # 2 and phantom #3 are shown in *Figure 7*. The linear relationship observed in phantom #1 was not reproduced, and no correlation with O_2_ concentration was observed in either case (R^2^ = 0.12 for phantom #2 and 0.04 for phantom #3). The slope of the linear fit in both phantoms was close to zero, as opposed to the linear and positive relationship that might have been expected from dissolved paramagnetic O_2_ molecules.

**Figure 7:**
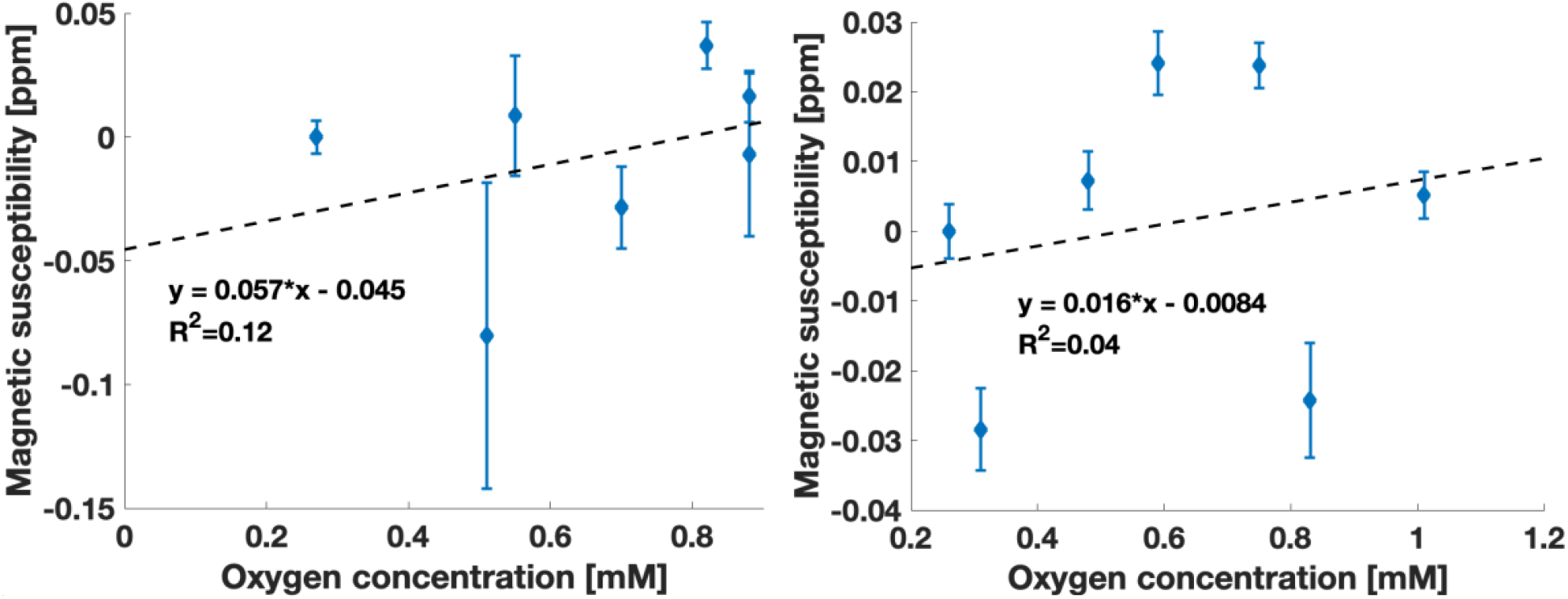
Variations of the magnetic susceptibility as a function of the *O*_2_ concentration measured in phantom #2 (left) and phantom #3 Exp. 1 with QSM (right). The mean (± standard deviation) magnetic susceptibility measured in 2D ROIs is plotted as a function of the measured *O*_2_ concentration. The dashed line corresponds to the linear fit. The vial at *O*_2_ concentration under atmospheric pressure (nominal *O*_2_ concentration of 0.29 mM) was used as the QSM reference.

Measurements of magnetic susceptibility in phantom # 3 using MR-susceptometry also did not reveal any trends with O_2_ concentration. The results obtained are shown in *Figure 8*. No correlation with O_2_ concentration was observed (R^2^ = 0.14). The susceptibility was nearly constant, and the slope of the linear fit was essentially zero.

**Figure 8:**
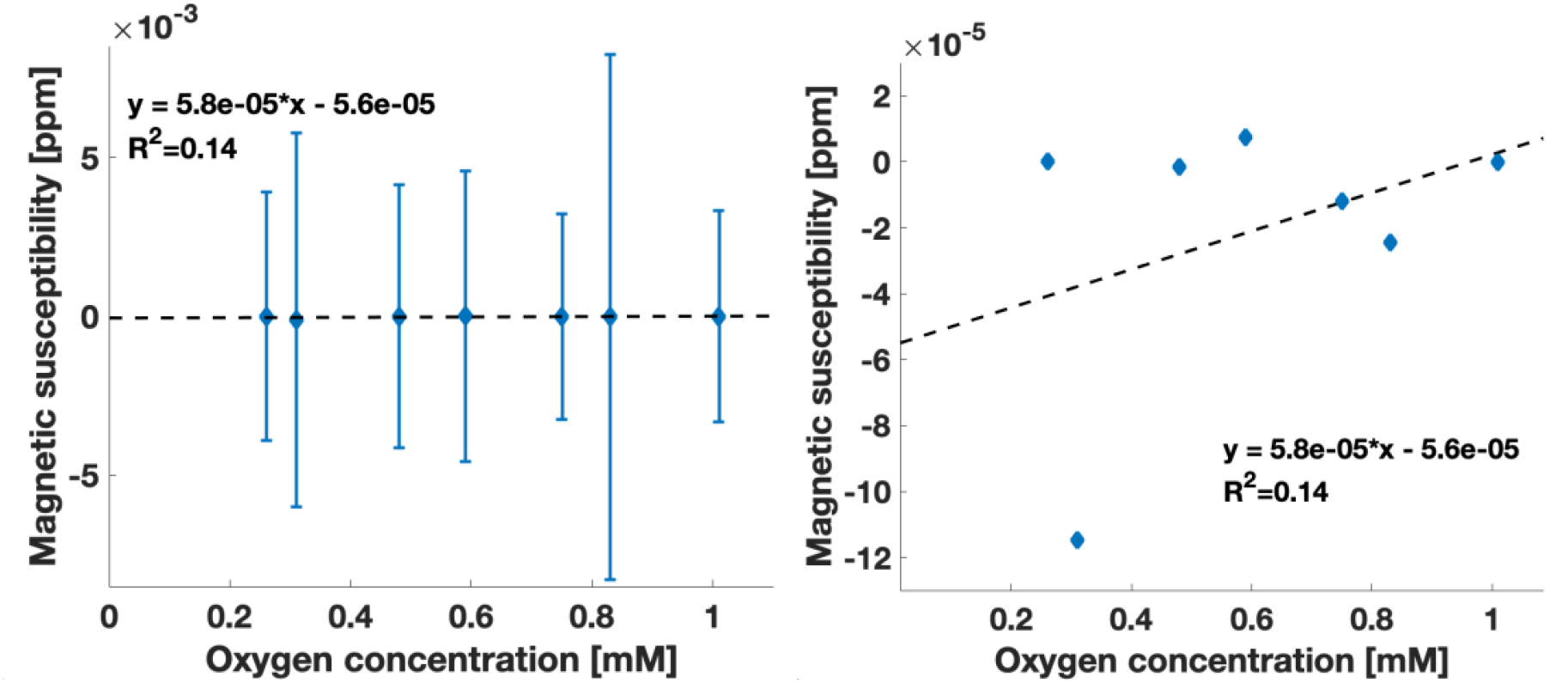
Variations of the magnetic susceptibility measured with MR susceptometry as a function of the *O*_2_ concentration measured in phantom #3 (Exp. 1). The dashed line corresponds to the linear fit. The vial at *O*_2_ concentration under atmospheric pressure (nominal *O*_2_ concentration of 0.29 mM) was used as the magnetic susceptibility reference. A zoomed in view is shown on the right panel without the errorbars to explain the non-zero coefficient of determination.

QSM measurements with long TEs in phantom #4 also did not reveal any trend for susceptibility as a function of O_2_ concentration. This is shown in *Figure 9*. Again, no correlation was observed between magnetic susceptibility and O_2_ concentration. The two points with N_2_ bubbling showed a more paramagnetic susceptibility compared to the point at O_2_ concentration under atmospheric pressure, which was unexpected, and matched observations in phantom #1. MR susceptometry measurements with long TEs in phantom #3 led to similar results, with no correlation (*Figure 10*). Two vials (#4 and #7) were removed from the analysis in this case due to the presence of an air bubble.

**Figure 9:**
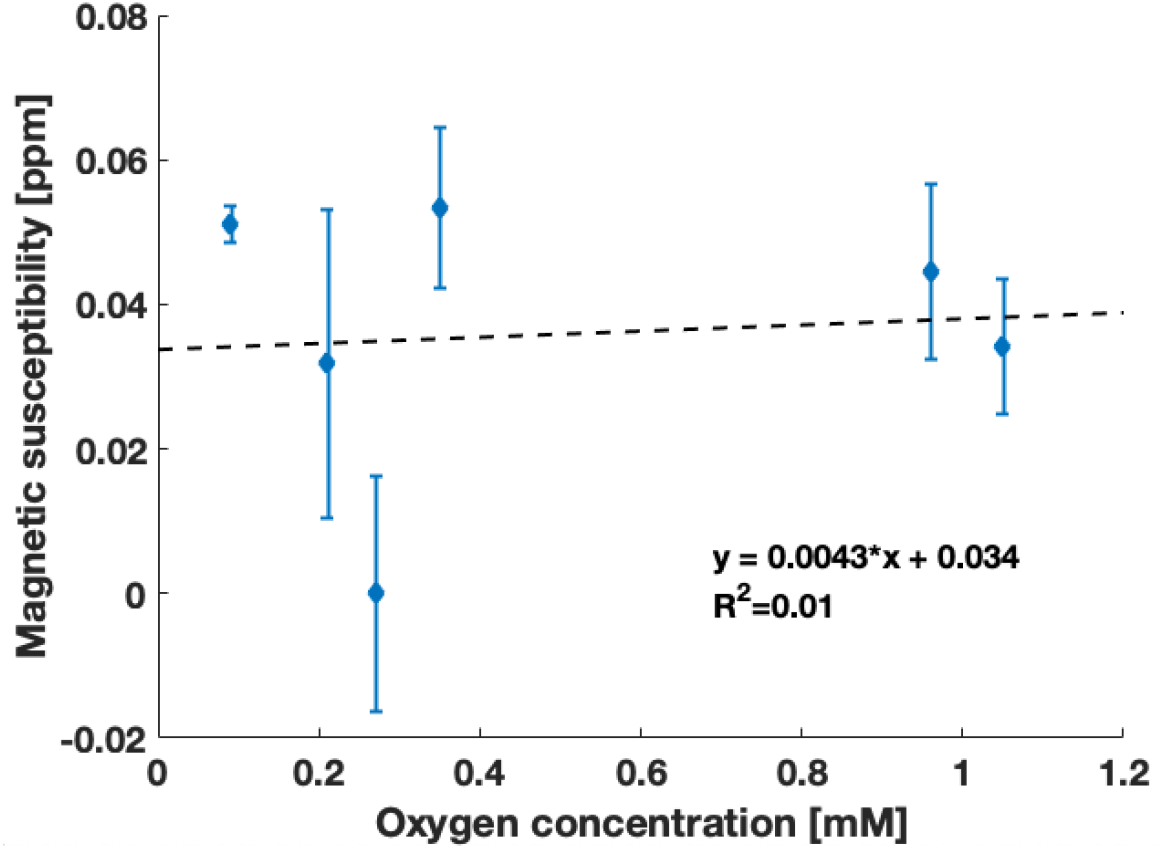
Variations of the magnetic susceptibility as a function of the *O*_2_ concentration measured in phantom #4. The mean (± standard deviation) magnetic susceptibility measured in a 2D ROI in each vial as a function of the measured *O*_2_ concentration is plotted. The dashed line corresponds to the linear fit. The vial at *O*_2_ concentration under atmospheric pressure (nominal *O*_2_ concentration of 0.29 mM) was used as the QSM reference.

**Figure 10:**
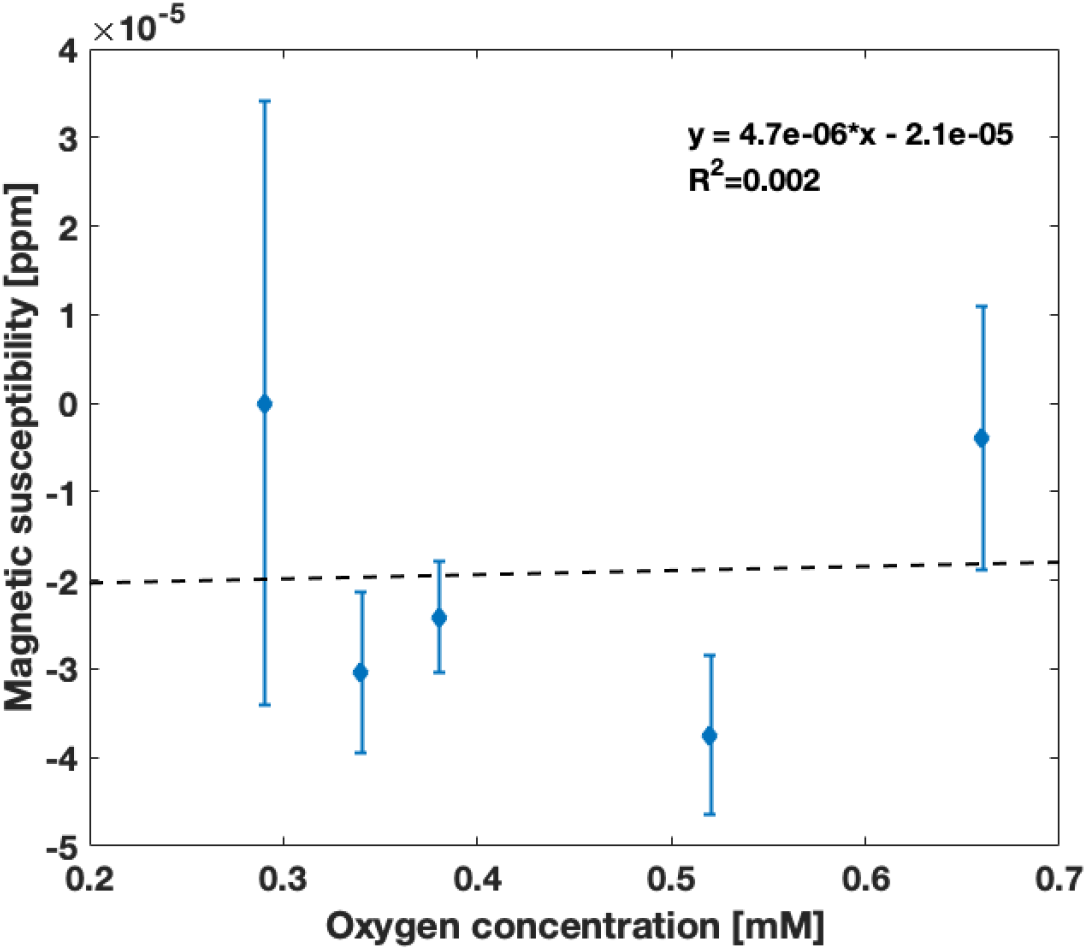
Variations of the magnetic susceptibility measured with MR susceptometry as a function of the *O*_2_ concentration measured in phantom #3 (Exp. 2) with long TEs. The dashed line corresponds to the linear fit. The vial at *O*_2_ concentration under atmospheric pressure (nominal *O*_2_ concentration of 0.29 mM) was used as the magnetic susceptibility reference.

## Discussion

This study hypothesized that QSM might be sensitive to variations in dissolved O_2_ concentration in fat-water mixtures. The hypothesis was tested in a set of experiments in dairy cream phantoms where the dissolved O_2_ concentration was controlled. These phantoms were envisioned as a surrogate material for human tissue, such that the experiments might mimic variations in tissue O_2_ concentration. Overall, the magnetic susceptibility measured with QSM clearly did not depend on the measured concentration of dissolved O_2_. This suggests that current QSM techniques are not precise enough to capture the O_2_ dependence of magnetic susceptibility.

The O_2_ concentration in dairy cream with high fat content was relatively stable over time, but it decreased over time in dairy products with lower fat content despite being placed in a sealed container. This observation might be explained by the consumption of O_2_ by the bacteria present in milk [32]. Light-induced oxidation could also be responsible for the observed decrease in O_2_ concentration as a function of time [33]. Therefore, 35% cream was the only good candidate for MR-oximetry phantoms, while milk and lower fat creams were not. Dairy products have been used in the literature as phantoms for T_2_ mapping [34], diffusion MRI [35], 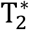 mapping [36], and T_1_ mapping [37]. However, there have been no reports in the literature on the use of dairy products as MR-oximetry phantoms. Phantoms based on isopropyl myristate oil and water emulsion have been proposed in the literature as liquid MR-oximetry phantoms [38]. Those emulsion phantoms require a complex preparation method. In contrast, dairy products are easy to obtain and highly controlled in terms of fat fraction content, making them an interesting alternative as MR-oximetry phantoms.

The use of N_2_ gas bubbling appeared to be a potential confounding factor for the investigation of the relationship between the magnetic susceptibility and O_2_ concentration. N_2_ gas bubbling resulted in a measurable decrease in dissolved O_2_ concentration, likely by replacement. A corresponding decrease in susceptibility would have been expected, as N_2_ gas is diamagnetic [23]. However, a positive shift in the magnetic susceptibility of vials bubbled with N_2_ was observed in phantoms #1 and #4 compared to the vial at O_2_ concentration under atmospheric pressure. This effect remains unexplained.

The results obtained in dairy cream phantoms with O_2_ gas bubbling suggest that QSM is not sensitive to dissolved O_2_ in liquid solutions. The absence of correlation between the magnetic susceptibility measurements performed with QSM in this work and dissolved O_2_ concentration contradicts a recent study performed with MR-susceptometry [5]. In that study, magnetic susceptibility showed a small but significant sensitivity to dissolved O_2_ in separate experiments in phantoms of water and blood plasma [5]. This could be due to differences between the techniques used: QSM versus MR susceptometry. MR susceptometry is potentially more accurate for MR-based measurements of magnetic susceptibility; however, it is not applicable in vivo because of its underlying geometry assumption. That is why QSM was the main focus of this work. The results obtained in this work suggest that current QSM techniques are not sensitive to variations in dissolved O_2_, perhaps due to measurement uncertainties. MR susceptometry was also evaluated in this work using phantom #3, but the results did not reveal any discernible trend of susceptibility with O_2_, as opposed to the previous report using susceptometry [5]. The lack of O_2_ sensitivity of the magnetic susceptibility measured in this work with MR-susceptometry might be also due to the short TEs that were initially used. Short TEs were selected to enable accurate fat-water separation, but this might have led to a lack of phase contrast accumulation over time. Additional MR susceptometry experiments were performed with additional longer TEs, on the order of 60 ms [5], but did not show any correlation between magnetic susceptibility and O_2_.

Our conclusion clashes with a recent study where QSM was performed in vivo in patients with brain tumors [15], in which a small increase in the magnetic susceptibility of tissues with low perfusion in response to a hyperoxic gas challenge was hypothesized to be associated with a change in dissolved O_2_ [15]. Based on the results obtained in our work, it appears unlikely that the change in magnetic susceptibility observed in vivo in that study is due to a change in dissolved O_2_ content. The reported small increase in magnetic susceptibility could be due not to change in tissue magnetic susceptibility in the tumor, but to a small decrease in magnetic susceptibility of the white matter region used as the reference region in that work. An increase in the O_2_ saturation state of blood hemoglobin, caused by the hyperoxic gas challenge, could have caused a decrease in susceptibility in the reference white matter region. This would artefactually appear as an increase in the magnetic susceptibility of the low perfused region after QSM referencing.

The lack of correlation between O_2_ concentration and QSM-based magnetic susceptibility measurements observed in this work could also be explained by the fact that measurements were performed in simple phantoms. QSM relies on MR signal phase variations introduced by various structures. It is challenging to build a phantom that contains enough structures to mimic in vivo anatomy. The lack of phase contrast in the phantoms used in this work could potentially lead to QSM inaccuracies. Future work could investigate the sensitivity of QSM to dissolved O_2_ directly in vivo in a controlled environment using oxygen electrodes inserted in tissues for O_2_ measurements, bypassing phantom experiments. O_2_ concentrations in vivo in tissues (ex: muscles) can be modulated by using hyperoxic gas challenges [39]. This approach could be used in future QSM experiments to investigate the potential of magnetic susceptibility as a biomarker for tissue oxygenation.

## Conclusion

To conclude, this work investigated the sensitivity of QSM-based magnetic susceptibility measurements to dissolved O_2_ in fat-water emulsions, as a model for lipid-containing tissues. The results obtained in phantoms were highly variable. The expected linear relationship between magnetic susceptibility and O_2_ concentration was not observed. This work suggests that the variations in magnetic susceptibility induced by the presence of dissolved O_2_ molecules are too small to be measured with QSM, given uncertainties of the current QSM techniques.

## Acknowledgments

The authors acknowledge the Body Magnetic Resonance research group (Technical University of Munich) for sharing their MRS processing software; the Cornell MRI Research lab (MEDI toolbox and online software) for making their QSM toolboxes available; Bruce Pike (University of Calgary) for lending the oxygen probe; Guillaume Gilbert (Philips Healthcare, Inc.), Avery Berman (Harvard University), Zaki Ahmed, and Stella Xing (McGill University) for useful discussion. This work was funded by the Montreal General Hospital Foundation, the Research Institute of the McGill University Health Centre, and a Discovery Grant from the Natural Science and Engineering Research Council of Canada (NSERC). VF acknowledges fellowship support from NSERC, and partial support from the NSERC CREATE Medical Physics Research Training Network (Grant number: 432290).

## Data availability statement

The code used for phase unwrapping with a quality-guided algorithm is available at: https://gitlab.com/veronique_fortier/Quality_guided_unwrapping. The code used for LBV and RESHARP background removal, and MEDI QSM reconstruction is available as part of the MEDI toolbox [40]: http://pre.weill.cornell.edu/mri/pages/qsm.html. Finally, the code used for 3-Point Dixon and IDEAL fat water separation is part of the ISMRM fat-water separation toolbox: https://www.ismrm.org/workshops/FatWater12/data.htm.

## Abbreviations used

Gd: Gadolinium
Hct: Hematocrit
IDEAL: Iterative Decomposition of Water and Fat with Echo Asymmetry and Least Squares Estimation
LBV: Laplacian Boundary Value
MEDI: Morphology Enabled Dipole Inversion MGRE Multi-Echo Gradient Echo
OEF: Oxygen Extraction Fraction
QBOLD: Quantitative Blood-Oxygen-Level Dependent QG Quality-Guided
QSM: Quantitative Susceptibility Mapping
RESHARP: Regularized-Enabled Sophisticated Harmonic Artifact Reduction for Phase data
ROI: Region Of Interest
STEAM: Stimulated Echo Acquisition Mode

