## Supporting information for "Dissolved oxygen minimally affects magnetic susceptibility in biologically relevant conditions"

### Characterization of the dissolved $O_2$ probe uncertainty

The dissolved $O_2$ probe was characterized over repeated measurements to evaluate its absolute uncertainty. Eleven 50 ml vials were filled with distilled water at room temperature and under $O_2$ concentration under atmospheric pressure. Three $O_2$ measurements were performed in each vial. The temperature was recorded for each measurement.

The probe yielded repeatable measurements within and between vials. The measured $O_2$ concentrations are shown in *Supporting Table S1*. An overall standard deviation of 0.24 mg/L was obtained over the 33 $O_2$ measurements. This corresponds to an absolute uncertainty of 0.01 mM, based on *Eq. 4*. The temperature decreased from 22.8 to 21.8 °C during the entire experiment.

### Characterization of the dissolved $O_2$ probe in water and dairy cream

The dissolved $O_2$ probe was characterized in pure distilled water and dairy cream with 35% of fat at room temperature. Ten 50 ml vials were filled with 42.5 ml of distilled water and six were filled with 42.5 ml of dairy cream. The vials were sealed with screw caps with a self-sealing cap membrane (SeptaSecure Uncut Cap, Syringa Lab Supplies Inc.), and thread seal tape was used to improve the seal of the screw cap. $O_2$ was added to the solutions by bubbling each vial with $O_2$ for different periods of time using a needle inserted through the membrane, connected to a controlled $O_2$ supply with a plastic tube. A second needle was inserted for air to flow out. An oxygen flow rate of 0.5 to 1 L/min was used.

The $O_2$ concentration and the temperature were measured in each vial using the dissolved $O_2$ meter. The temperature varied from 20 to 22 °C. The results of the $O_2$ measurements as a function of the $O_2$ bubbling time are illustrated in *Supporting Figure S1*.

Both water and dairy cream were saturated to an $O_2$ concentration of approximately 40 mg/L. This is slightly below the full saturation level of 45 mg/L, which could likely not be reached because of the experimental setup, where there was a small air volume at the top of each solution in all vials. The fact that both solutions saturated to the same $O_2$ concentration demonstrates the validity of the $O_2$ measurements in dairy cream.

**Supporting information figure**

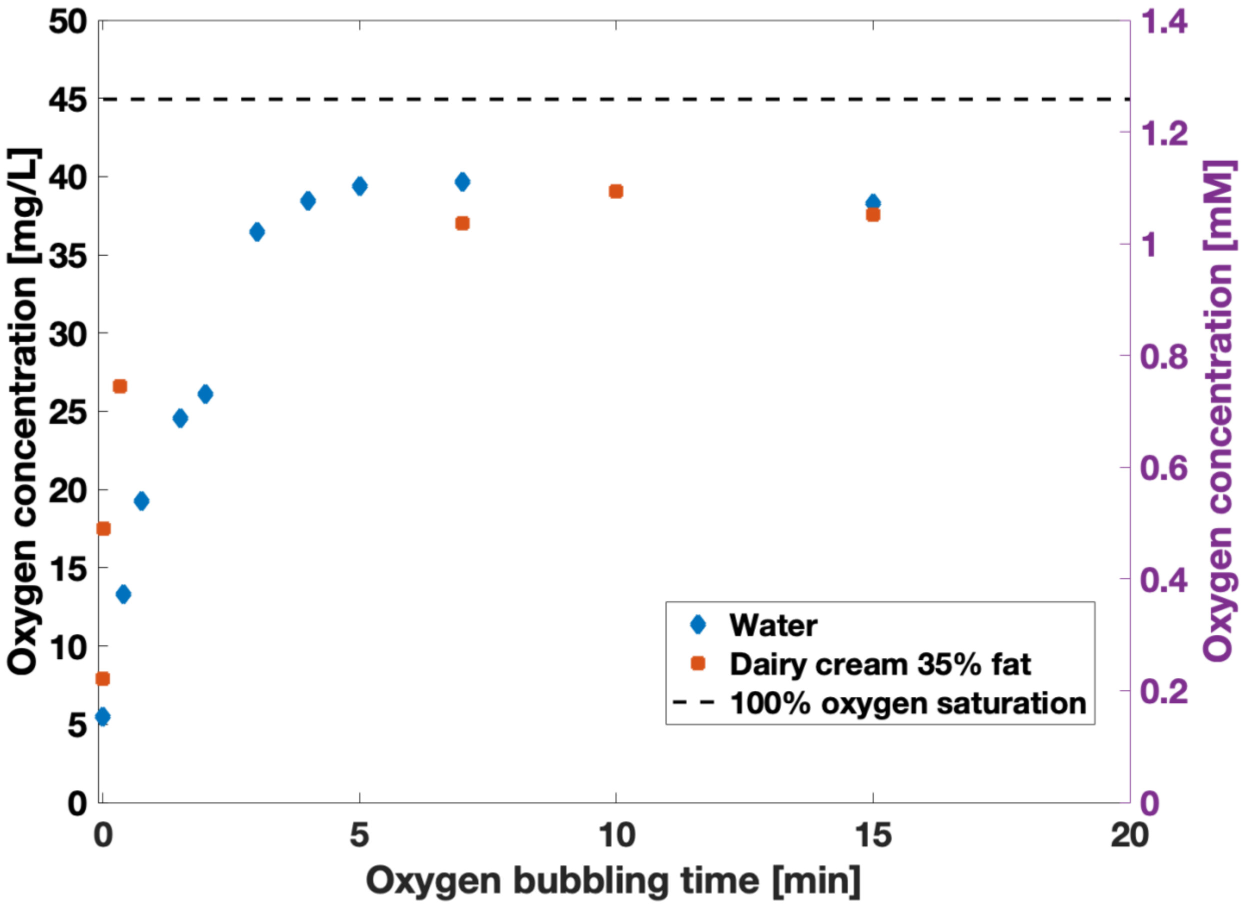


Supporting Figure S1: $O_2$ concentration as a function of $O_2$ bubbling time in distilled water and dairy cream with 35% of fat.

**Supporting information table**

Supporting Table S1: $O_2$ concentration measured in all vials over three repeated measurements. The mean and standard deviation is also included for each vial, as well as for all 33 measurements combined.

| | Oxygen concentration (mg/L) | | | | |
|---|---|---|---|---|---|
| **Vial** | **#1** | **#2** | **#3** | **Mean** | **Std** |
| **1** | 6.59 | 6.67 | 6.81 | 6.69 | 0.11 |
| **2** | 6.96 | 7.05 | 7.05 | 7.02 | 0.05 |
| **3** | 6.79 | 7.02 | 7.1 | 6.97 | 0.16 |
| **4** | 6.65 | 6.87 | 6.94 | 6.82 | 0.15 |
| **5** | 7.05 | 6.75 | 6.89 | 6.90 | 0.15 |
| **6** | 7.13 | 6.98 | 7.02 | 7.04 | 0.08 |
| **7** | 7.15 | 7.14 | 7.26 | 7.18 | 0.07 |
| **8** | 7.39 | 6.93 | 7.28 | 7.20 | 0.24 |
| **9** | 6.96 | 7.04 | 6.84 | 6.95 | 0.10 |
| **10** | 7.09 | 7.24 | 6.97 | 7.10 | 0.14 |
| **11** | 7.1 | 7.21 | 7.34 | 7.22 | 0.12 |
| **Combined** | - | - | - | 6.99 | 0.24 |
